# Phyling: phylogenetic inference from annotated genomes

**DOI:** 10.1101/2025.07.30.666921

**Authors:** Cheng-Hung Tsai, Jason E. Stajich

## Abstract

Phyling is a fast, scalable, and user-friendly tool supporting phylogenomic reconstruction of species phylogenies directly from protein-encoded genomic data. It identifies orthologous genes by searching a sample’s protein sequences against a Hidden Markov Models marker set, containing single-copy orthologs, retrieved from the BUSCO database. In the final step, users can choose between consensus and concatenation strategies to construct the species tree from the aligned orthologs.

Phyling efficiently resolves large phylogenies by optimizing memory usage and data processing. Its checkpoint system enables users to incrementally add or remove samples without repeating the entire search process. For analyses involving closely related taxa, Phyling supports the use of nucleotide coding sequences, which may capture phylogenetic signals missed by protein sequences. The benchmark results show that Phyling substantially runs faster than OrthoFinder, a Reciprocal Best Hit based method, while achieving equal or better accuracy.

## Introduction

Phylogenetic analysis provides the framework for understanding the evolutionary relationships among organisms, which is fundamental to modern biology. Phylogenetic trees built from genetic data can trace the origin and evolution of genes, species, and traits; identify conserved biological functions; and reveal the processes driving species diversification and adaptation. (Soltis and Soltis 2003; Yang and Rannala 2012)

Phylogenetic inference depends on models of sequence divergence, so choosing the right sequences is crucial to accuracy. Orthologous sequences, those that diverged through speciation from a common ancestor, are typically the most appropriate markers for reconstructing species relationships. (Kapli *et al*. 2020) A major strategy for identifying orthologs is similarity-based comparison, which clusters genes with high sequence similarity. (Tekaia 2016; de Boissier and Habermann 2020) Two common implementations of this approach are Reciprocal Best Hit (RBH) and profile-based matching.

RBH identifies orthologs by finding genes in two genomes that are each other’s best match in bidirectional sequence similarity searches. In this approach, an all-against-all search is applied across samples, and clusters of RBH pairs are identified as orthologs. Several tools, such as OrthoMCL (Li *et al*. 2003), OrthoFinder (Emms and Kelly 2019), and OrthoPhy (Watanabe *et al*. 2023) adopt RBH combined with other clustering techniques like Markov Clustering to infer orthology.

In contrast, Profile-based sequence matching identifies orthologs by clustering the sequences that match to the same profile in a database with known orthologous groups. Users match sequences in a genome against databases such as eggNOG (Hernández-Plaza *et al*. 2023), TreeFam (Ruan *et al*. 2008), and Pfam (Paysan-Lafosse *et al*. 2025), to classify the input sequences based on orthology or function. Though conceptually distinct, RBH and profile-based matching are often used together. For example, eggNOG uses RBH for initial ortholog detection and profile Hidden Markov Models (HMMs) for efficient large-scale inference. This hybrid approach balances RBH’s precision with the scalability of profile-based methods for better phylogenetic inference.

While all-against-all searches usually offer higher accuracy, they become computationally expensive as more genomes are added due to the quadratic growth in pairwise comparisons. Moreover, the rigid structure of the RBH-based pipelines make it less flexible when updates are needed. Any change, such as adding or removing a single genome, requires repeating the entire search and clustering process. In contrast, profile-based approaches allow each sample to be processed independently, so users can incrementally update datasets without rerunning previously completed searches. This makes profile-based methods particularly convenient for large or frequently updated phylogenomic datasets.

Here we present Phyling, a fast and user-friendly tool for inferring phylogeny directly from genome data. Phyling utilized the profile-based ortholog identification strategy to achieve significant gains in speed and resource efficiency over RBH-based methods. Additionally, it can handle large-scale datasets of up to thousands of species, which are usually challenging for other tools. Phyling also features a checkpoint design that allows users to efficiently redo analyses without re-run searches on previously processed samples, further enhancing its scalability.

## Materials and methods

### Phyling pipeline

Phyling was mainly developed in Python >= 3.9 and relies on a minimal set of command-line tools. To simplify installation and prevent dependency conflicts, we use Conda for managing the software environment and installing all required dependencies. The Python package Biopython >= 1.81 (Cock *et al*. 2009) was used for sequence data processing. To allow more flexibility and user customization, Phyling is organized into four modules - **download, align, filter** and **tree** (Fig. 1).

**Figure 1.**
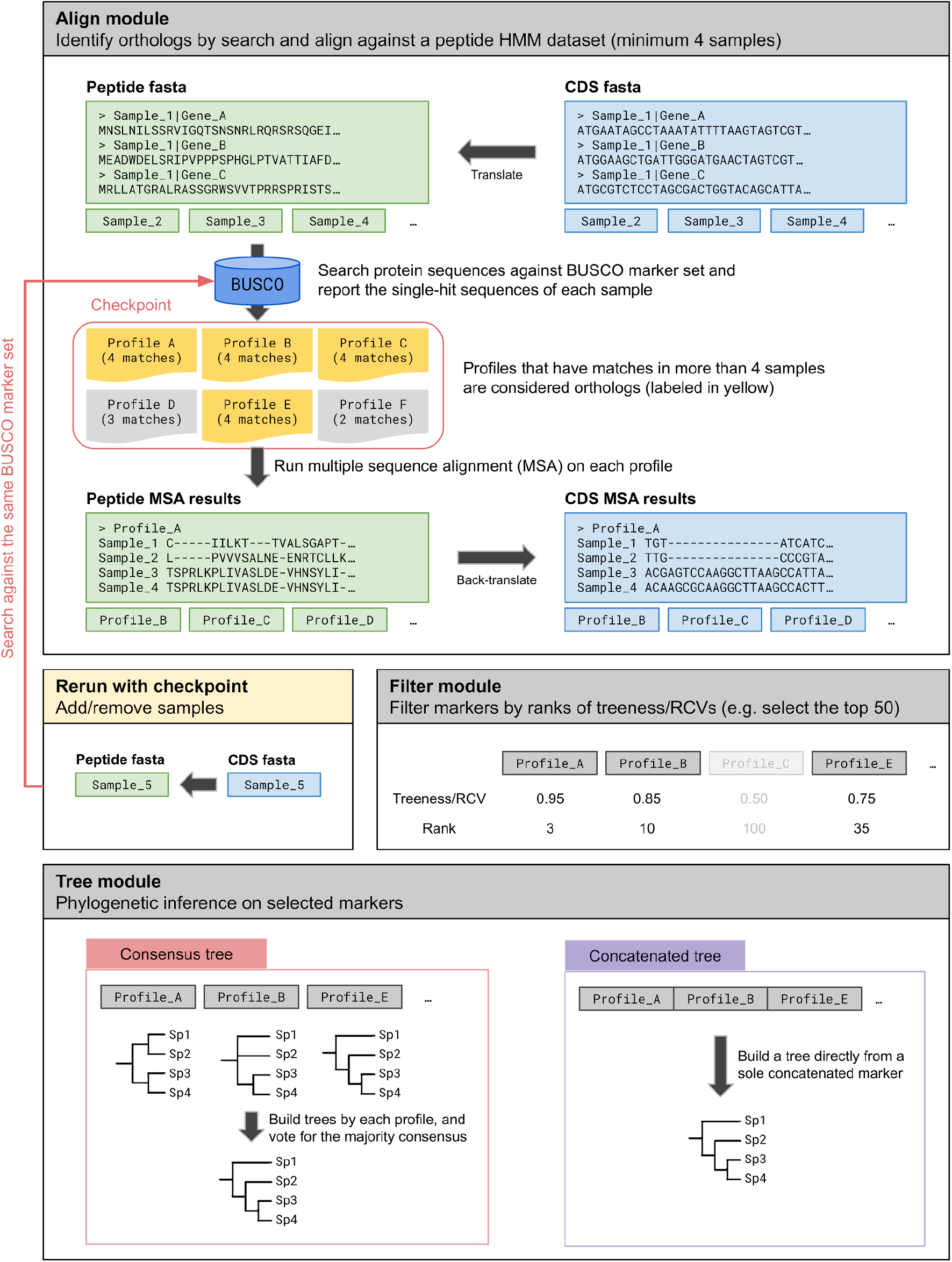
Flowchart of Phyling’s core modules - align, filter and tree. The download module is not shown. The salmon-colored arrow indicates the checkpoint mechanism, in which hits from a newly added sample are incorporated into the existing hits collection from previous runs.

The **download** module facilitates the retrieval of HMM profiles for the selected marker sets from the BUSCO database (Manni *et al*. 2021), which are used to identify orthologs in the input data. Upon first use, Phyling creates a directory under the user’s home folder to store the downloaded marker sets.

The **align** module identifies the orthologs from each sample and further aligns their sequences. Phyling accepts either proteomes or their nucleotide coding sequences with a minimum of four samples. The sequences of each sample are searched in parallel against the HMM profiles of a marker set using hmmsearch from pyHMMER v0.11.0 (Larralde and Zeller 2023). The searching parameters as well as the search hits will be saved in a checkpoint file for future rerun. To ensure orthology, any sample with multiple hits to the same HMM profile was excluded from further analysis for that marker. The sequences corresponding to these hits, which match to the same HMM profile, are extracted using pyfaidx v0.8.1.3. (Shirley *et al*. 2015) By default, the extracted sequences are aligned using hmmalign from pyHMMER package. Users can switch to use Muscle v5.3 (Edgar 2022) to generate higher-quality alignments. The alignments will be trimmed by ClipKIT v2.1.1 (Steenwyk *et al*. 2020) to report only the parsimony-informative sites.

The **filter** module selects the most informative marker alignments for phylogenetic inference. Phyling uses FastTree v2.1.1 (Price *et al*. 2010) to construct trees for each marker. These trees are then evaluated using the treeness over relative composition variability (RCV) score, calculated via PhyKIT v2.0.1 (Steenwyk *et al*. 2021), to quantify their phylogenetic informativeness. Markers are ranked by their treeness/RCV scores, and the top n markers, as specified by the user, are retained for downstream analysis.

The **tree** module conducts the final phylogenetic inference using the filtered marker alignments. By default, Phyling applies a consensus strategy, constructing individual gene trees for each marker and inferring a species tree based on their congruence using ASTER v1.19 (Zhang *et al*. 2018), a C-based implementation of ASTRAL. Alternatively, users may opt for a concatenation approach, where all marker alignments are combined into a single supermatrix. In the concatenation workflow, Phyling utilizes ModelFinder (Kalyaanamoorthy *et al*. 2017) from the IQ-TREE package to determine the best-fit substitution model. Users can select either FastTree, RAxML-NG (Kozlov *et al*. 2019), or IQ-TREE (Minh *et al*. 2020) for inference based on their requirements. For assessing branch support, both ultrafast bootstrap (UFBoot) (Hoang *et al*. 2018) and site concordance factor (Mo *et al*. 2023) are employed. When using RAxML-NG or IQ-TREE in concatenation mode, users can optionally enable partitioned analyses, which assign distinct substitution models to each marker (Chernomor *et al*. 2016), allowing more accurate modeling of heterogeneity across loci.

### Dataset for benchmark

To demonstrate the performance of Phyling, we employed both Ensembl Genomes 60 datasets (Yates *et al*. 2022) and a simulated dataset for the benchmark. For Ensembl datasets, the metadata for 31,332 bacterial strains and 1,505 fungal strains were downloaded and then selectively subset for different benchmarking scenarios.

To evaluate Phyling’s performance under different phylogenetic scenarios, we subdivided the Ensembl datasets into two types. The first type, hereafter referred to as the general dataset, mimics more common use cases by focusing on samples within the same family. The fungal general dataset includes 28 samples from the family *Saccharomycetaceae* (class *Saccharomycetes*), with an additional two samples from the family *Phaffomycetaceae* included as outgroup taxa, resulting in a total of 30 fungal samples. The bacterial general dataset comprises 96 samples - 94 from the family *Staphylococcaceae* and two outgroups from *Bacillaceae* (order *Bacillales*). To simulate more challenging inference conditions involving greater evolutionary divergence, we also constructed distant datasets by selecting one representative strain per family and per order from the full fungal and bacterial sets. This resulted in 251 samples in the bacterial dataset and 231 in the fungal dataset.

The simulated dataset was obtained from the work of Lees et al. (Lees *et al*. 2018) which comprises 96 *Streptococcus pneumoniae* strains. The dataset, artificially evolved from the reference genome *S. pneumoniae* ATCC 700669 using a known phylogenetic framework originally inferred by Kremer et al. (Kremer *et al*. 2017), serve as a ground-truth for benchmarking the accuracy of phylogenetic inference.

### Benchmark

To monitor computational resource usage, including time and memory consumption, we employed the Python package memory-profiler version 0.61.0 (Pedregosa 2022) during all benchmarks. Each tool was executed three times under identical conditions on a high-performance computing cluster equipped with AMD EPYC 7713 64-core processors (base clock 2GHz, boost clock up to 3.67 GHz) to ensure consistency and reproducibility of the performance measurements.

Generalized Robinson–Foulds (RF) distance implemented in R library TreeDist (Smith 2020) was used to measure distances between trees inferred by different workflows. Non-metric multidimensional scaling (NMDS) was performed on RF distance matrices using the Python package scikit-learn (Pedregosa *et al*. 2011) for visualization. The R library MonoPhy (Schwery and O’Meara 2016) was used to assess the monophyly status of tree pairs constructed by different workflows, as shown in Table S3 and Figures S3, S4 and S5.

### Data visualization

We use the Python package matplotlib and its high-level interface seaborn to visualize most of the results. (Hunter 2007; Waskom 2021) R libraries ggplot2 and ggtree were used for phylogenetic tree visualization. (Wickham 2009; Yu *et al*. 2017)

## Results

### Phyling runs with less resources but faster using comparable tree building algorithms

The optimized parallelization and data processing design contributes to the speed of Phyling and allows it to perform analysis on over thousands of samples. Overall, Phyling runs most efficiently with 16 threads when benchmarking with the fungal distant dataset and the acceleration has limited improvement when applying more than 32 threads. (Fig. 2) Therefore, 16-thread configuration will be used in the following comparison benchmarks.

**Figure 2.**
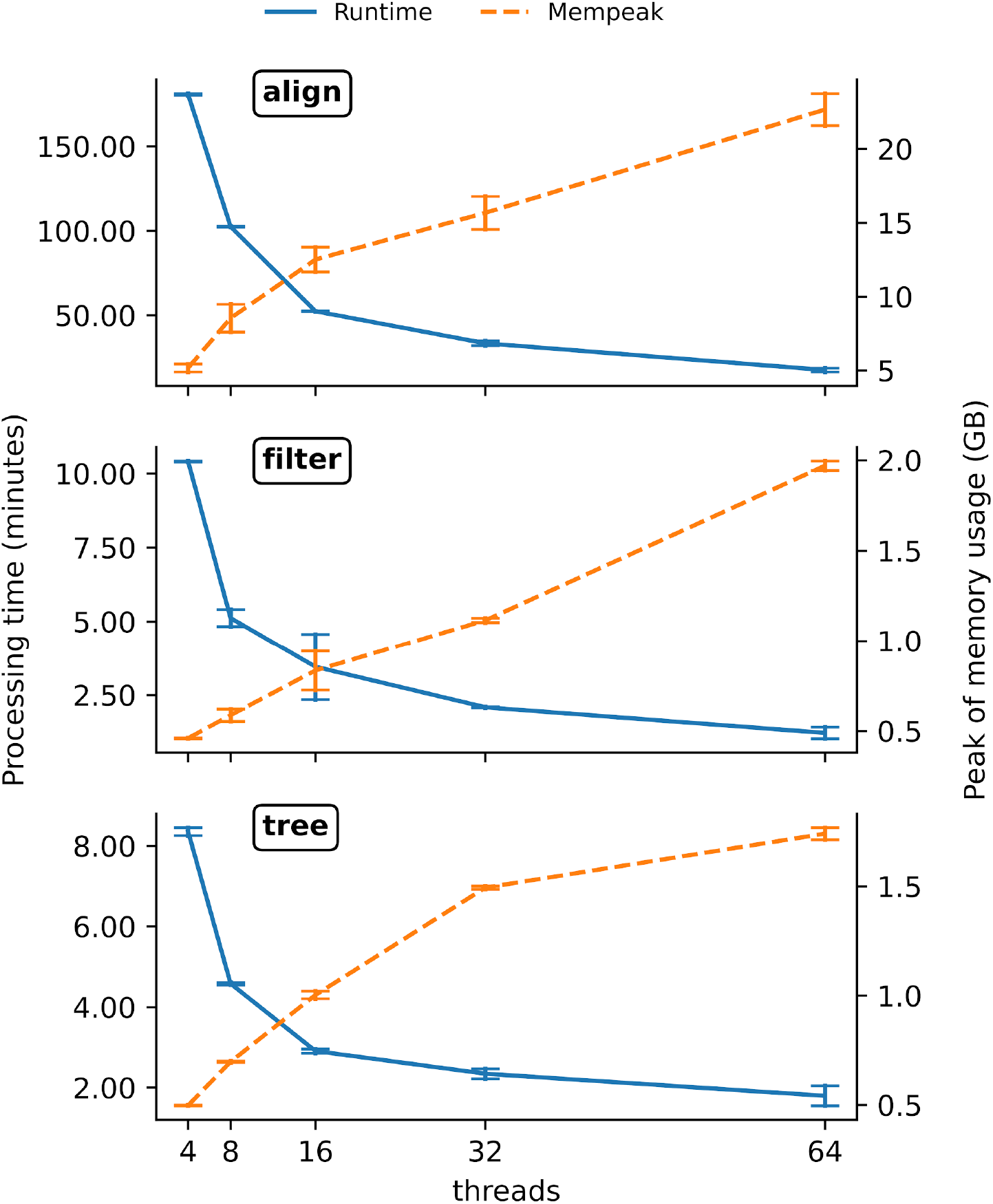
The multithreading performance benchmark for each module of Phyling. Blue solid lines (left y-axis) represent runtime in minutes and orange dashed lines (right y-axis) indicate peak memory usage in GB. Error bars represent standard deviations based on three replicates.

To benchmark the performance of Phyling, we compared it with OrthoFinder, one of the widely used phylogenetic inference tools based on RBH, and GToTree (Lee 2019), a tool which also developed upon profile-based approach to demonstrate the performance. When benchmarking the bacterial distant dataset with BUSCO bacteria_odb10 marker set (contains 124 markers), Phyling took less than five minutes on average with consensus mode, faster than both OrthoFinder and GToTree (using Bacteria SCG-set which contains 74 profiles) that took approximately 11 and 1 hour, respectively. In contrast, the more comprehensive concatenation mode took over eight hours which is still faster than OrthoFinder. (Fig. 3 and Table S1) The UFBoot accounted for a large portion of the time spent in concatenation mode. By default, GToTree runs FastTree to infer phylogenies from concatenated sequences. To enable a more direct comparison, we used the concatenated sequences generated by Phyling’s tree module and ran FastTree independently. With this alternative workflow, Phyling completed the analysis in approximately 10 minutes, faster than GToTree, which required nearly an hour. Phyling’s peak usage was only about 10-15% of that of OrthoFinder in the bacterial dataset depending on the inference mode. (Table S2) Both Phyling and GToTree are very memory efficient tools, using at most 5 GB during the run.

**Figure 3.**
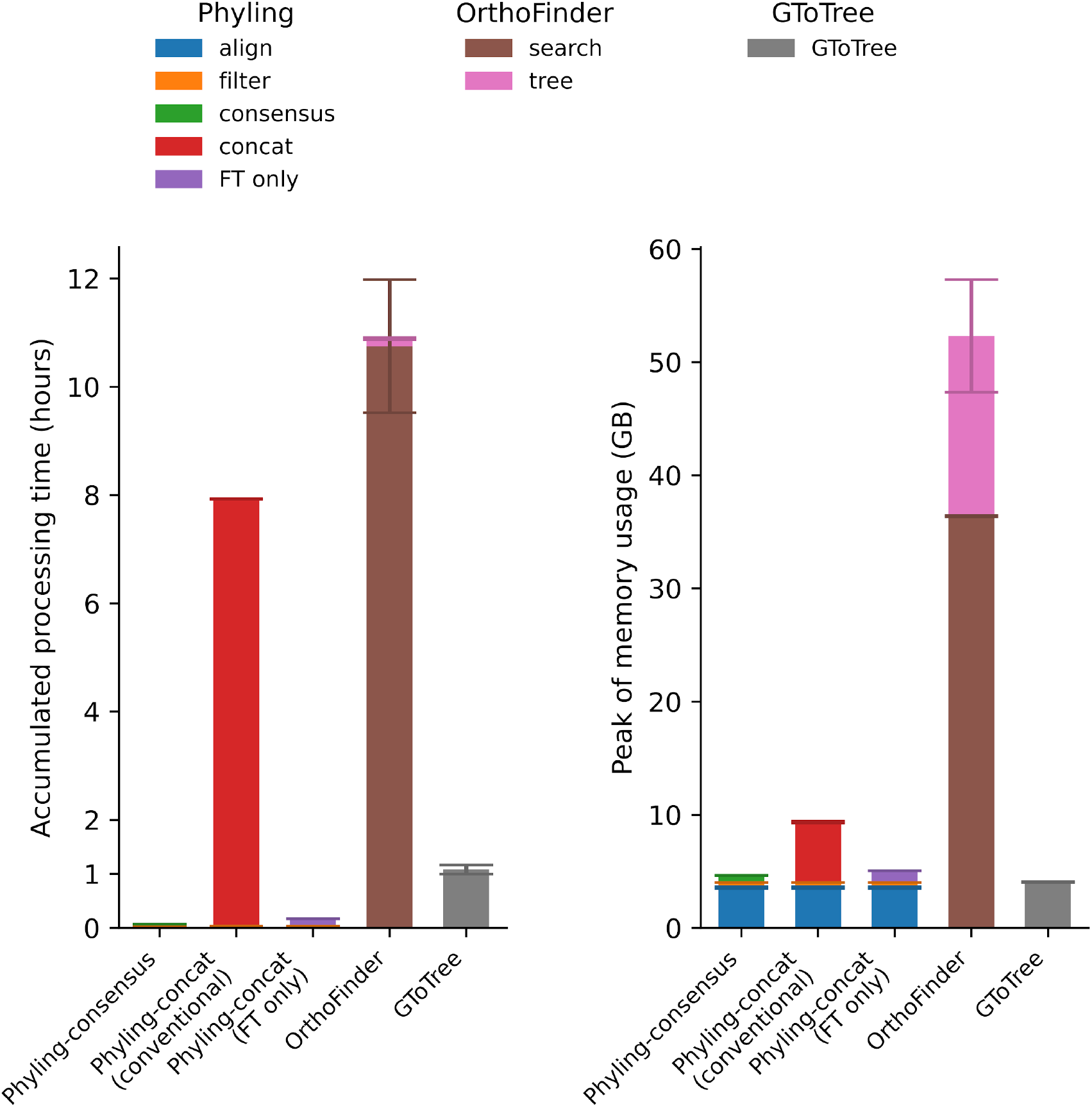
Accumulated runtime and peak of memory usage for different modes of Phyling, OrthoFinder and GToTree on bacterial distant dataset. The conventional concatenation mode of Phyling performs best-fit model selection, followed by tree inference with FastTree, and branch support assessment as described in Materials and Methods section. The alternative “FT only” concatenation mode runs FastTree with LG model and discrete gamma distribution directly on the concatenated sequence made by the conventional pipeline. Colors represent different tools and modules. Error bars represent standard deviations based on three replicates.

When benchmarking the fungal distant dataset with BUSCO fungi_odb10 marker set (contains 758 markers), Phyling completed the analysis in less than one hour on average using the consensus mode, and approximately 10 hours with concatenation mode. Both modes were substantially faster than OrthoFinder, which took around 45 hours to complete. (Fig. 4 and Table S1) The major factor contributing to the difference in processing time is the number of orthologs used for phylogenetic inference. OrthoFinder identified 91,553 orthogroups in its analysis, whereas Phyling utilizes only 754. We do not include the GToTree results since there is no suitable markerset for fungi inference. The only relevant markerset, Universal-Hug-et-al (Hug *et al*. 2016), contains only 16 markers and most of the genomes were removed during the filtering process. In terms of memory usage, Phyling’s peak memory consumption was only about 10% of that observed for OrthoFinder. (Table S2)

**Figure 4.**
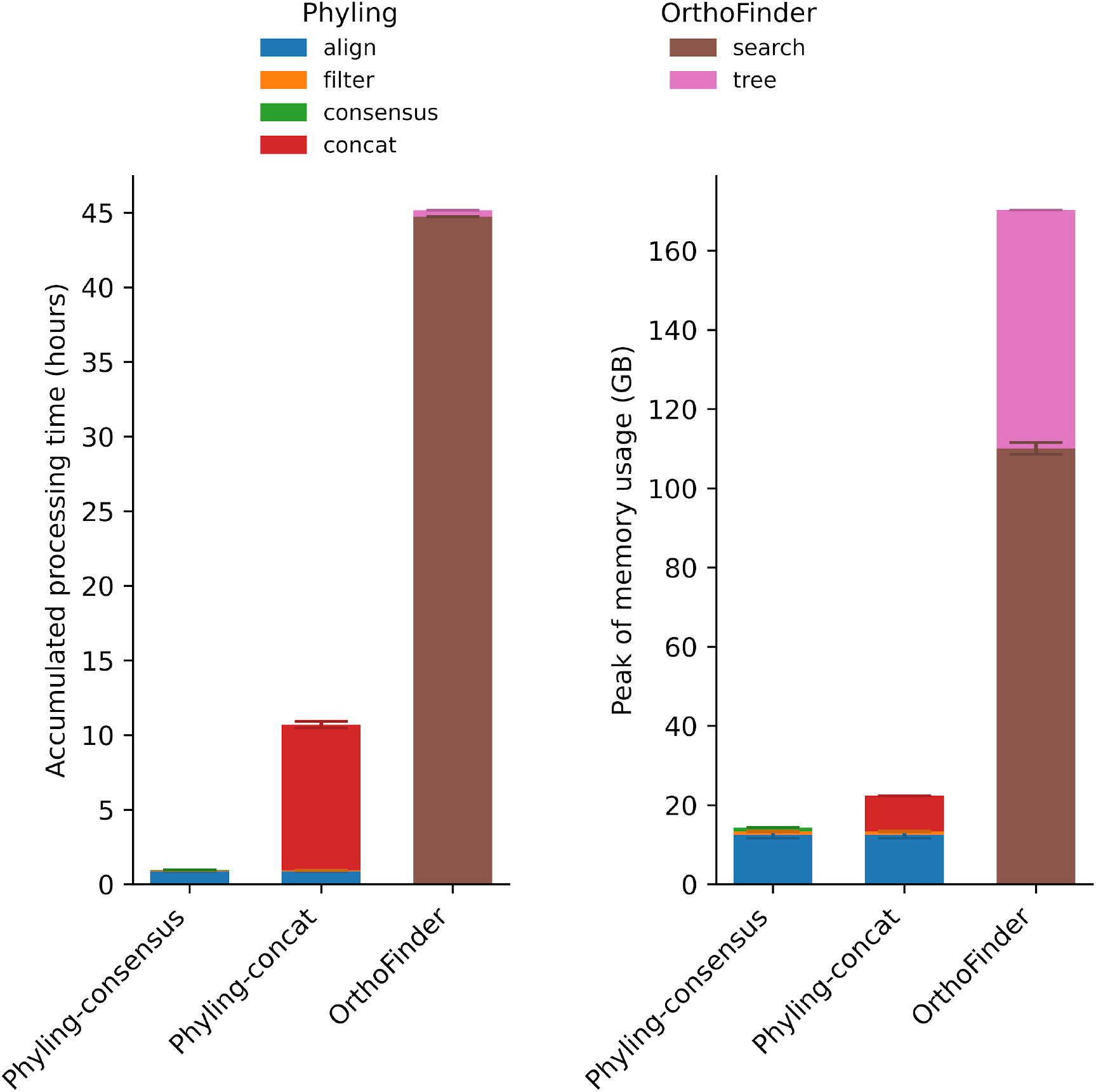
Accumulated processing time and peak of memory usage for different modes of Phyling, OrthoFinder and GToTree on fungal distant dataset. Colors represent different tools and modules. Error bars represent standard deviations based on three replicates.

### Phyling achieves equal or better accuracy

To assess the accuracy of Phyling’s phylogenetic inference, we computed the generalized RF distance between its trees and those constructed by OrthoFinder and GToTree across both the general and distant datasets. Overall, the inferences made by Phyling, under both consensus and concatenation modes, are highly congruent with those made by OrthoFinder and GToTree, in both bacterial and fungal general datasets. (Fig. 5 and Fig. S1) When evaluated using the fungal distant dataset, trees generated by Phyling also exhibited a high degree of topological similarity to those produced by OrthoFinder. (Fig. 6 and Fig. S2) This similarity between the trees was higher when all available marker genes were used rather than using a filtered subset restricted to the top 50 scoring markers. Benchmarking with the bacterial distant dataset revealed greater topological divergence among the three workflows, suggesting higher complexity or variability in bacterial phylogenetic signals. When using phylum-level taxonomy to assess the monophyly of resulting trees, all three methods performed comparably with GToTree having a slightly higher number of monophyletic taxa. (Fig. S3, S4 and S5)

**Figure 5.**
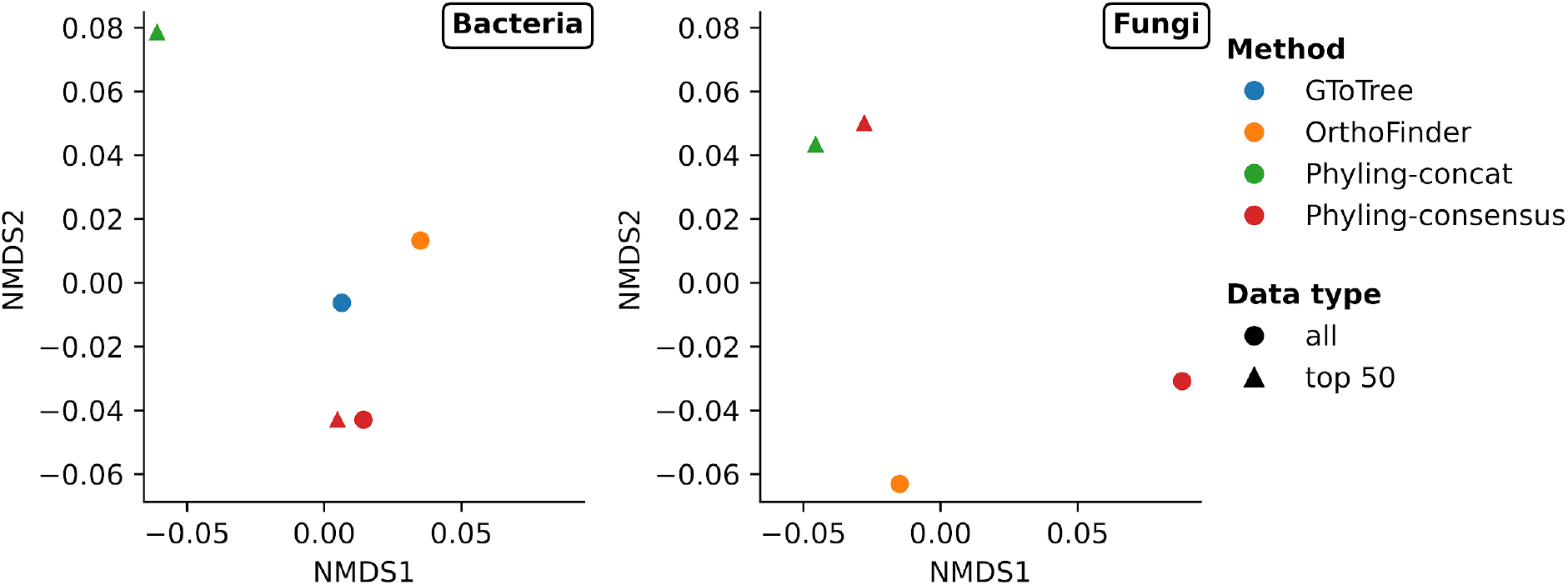
The NMDS ordination plot of RF distances among trees inferred from general datasets. The left and right plots represent results from the bacterial and fungal general datasets, respectively. Colors represent different methods and shapes for data types in NMDS plots.

**Figure 6.**
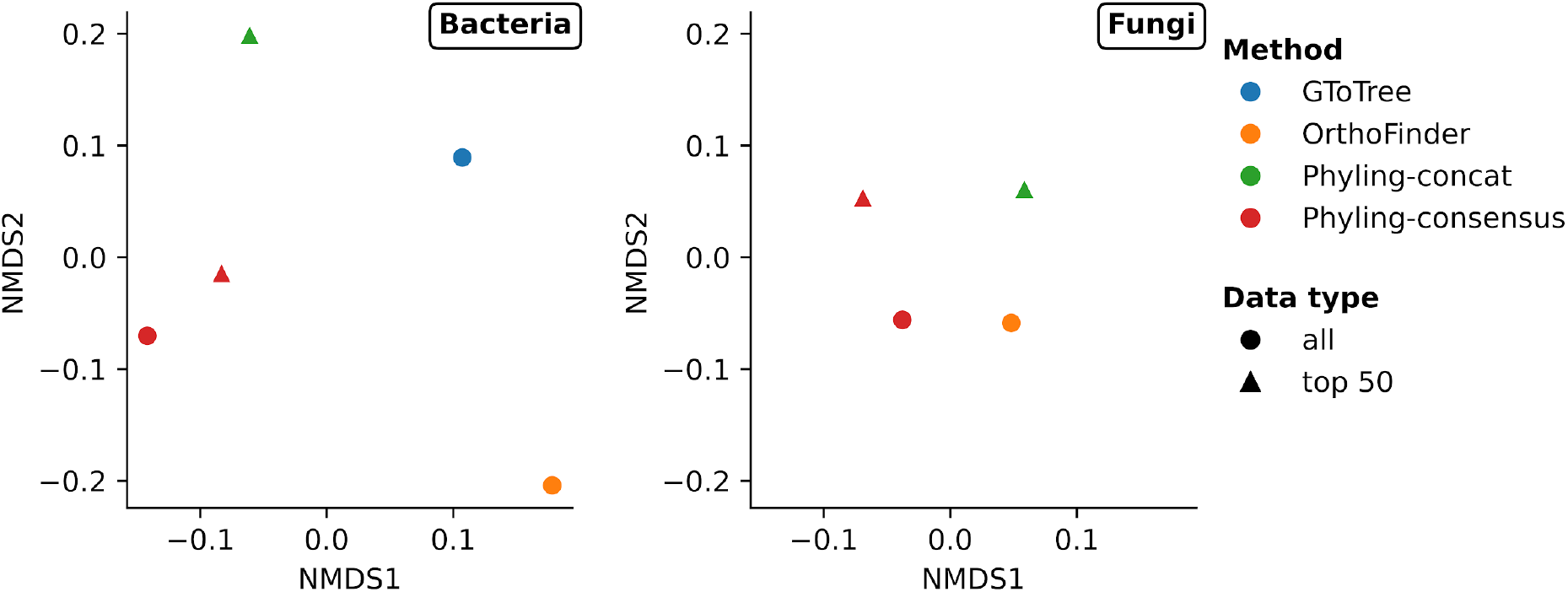
The NMDS ordination plot of RF distances among trees inferred from distant datasets. The left and right plots represent results from the bacterial and fungal distant datasets, respectively. Colors represent different methods and shapes for data types in NMDS plots.

To further assess Phyling’s performance on datasets comprising close lineages, we evaluated it using a simulated dataset with known phylogenies described by Lees et al.. Across all three workflows, phylogenetic inferences based on peptide sequences were generally far less accurate than those derived from coding nucleotide sequences. (Fig. 7) Among the workflows, the Phyling tree had the highest topological similarity to the reference tree when using coding nucleotide sequences, particularly when inferring by the concatenation mode. The accuracy slightly improved when using the streptococcaceae_odb12 marker set (contains 689 markers) under both the concatenation mode and the consensus mode utilizing all markers, compared to the more general bacteria_odb10 marker set. However, the phylogenetic tree inferred using only the top 50 markers from the streptococcaceae_odb12 set showed the lowest accuracy among all tested combinations of marker sets and inference modes. This result highlights that employing a taxon-specific marker set does not consistently guarantee optimal performance, particularly when using a reduced subset of markers. (Fig. S6)

**Figure 7.**
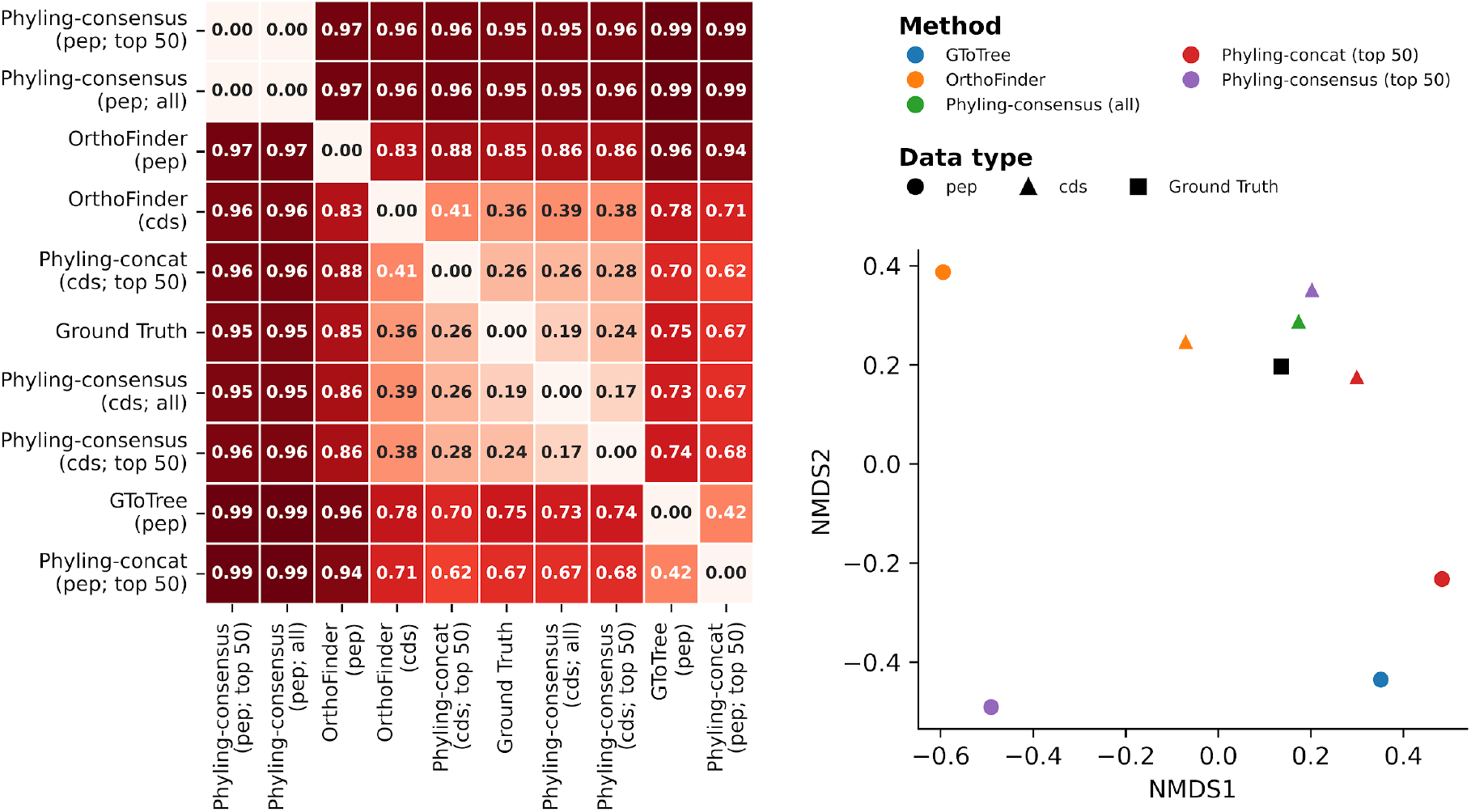
Generalized RF distance matrix and its corresponding NMDS plot among trees inferred from the simulated dataset. The distance matrix on the left is hierarchical clustered, with values ranging from 0 to 1. In the NMDS plot, colors represent different methods, shapes represent data types, and the black square indicates the ground truth. Nucleotide-based inference by GToTree was not performed, as this option is only available when working directly with unannotated genomes.

## Discussion

Phylogenetic analysis is a crucial step in comparative and population genomics, and Phyling streamlines this process with speed and accuracy. Taking advantage of the extensive marker collections from BUSCO, users can easily select an appropriate marker set for their analyses. Even with a more generalized marker set, Phyling still can offer relatively accurate results. (Fig. 7) By employing HMM-based profile matching and optimized parallelization strategy through PyHMMER, Phyling processes large datasets significantly faster than RBH-based methods and even traditional HMMER searches. With all these user-friendly setups and processing speed, Phyling does not sacrifice the accuracy of the inference.

An additional advantage of using the BUSCO marker set for ortholog identification is the reduction of errors in phylogenetic inference caused by gene duplication events. Since phylogenetic reconstruction fundamentally depends on accurately identified orthologs, misidentification can lead to incorrect tree topologies. (Tekaia 2016; Emms and Kelly 2019; de Boissier and Habermann 2020; Kapli *et al*. 2020; Steenwyk *et al*. 2023) The BUSCO marker sets are composed of genes that are expected to be present in single copy across a wide taxonomic range, thereby minimizing the inclusion of paralogs. Furthermore, Phyling incorporates an additional filtering step, similar to that implemented in GToTree (Lee 2019), to exclude sequences from a sample if multiple hits are found for the same marker to further enhance orthology.

The concatenation mode offers more dedicated inference and additional branch support information than speed-focus consensus mode. In addition to bootstrap values, Phyling incorporates site concordance factors from IQ-TREE to provide users with complementary insights into gene tree discordance. This is particularly valuable given that topological variation is not always captured by bootstrap values, especially when inferencing from a concatenated dataset. (Lanfear and Hahn 2024) However, these steps increase runtime. Automatic model selection and bootstrapping are the main contributors, as both involve iterative optimization. In particular, UFBoot may take longer to converge when samples are highly phylogenetically divergent.

Profile-based methods offer not only speed but also significant advantages in memory efficiency. In RBH-based approaches, identifying orthogroups requires exhaustive pairwise comparisons across all samples, which can create a bottleneck on systems with limited memory due to the accumulation of intermediate files. In contrast, the profile-based methods directly identify best hits against a predefined marker set without the need for pairwise comparisons, resulting in much lower memory usage. This efficiency enables Phyling to scale effectively to datasets with thousands of samples.

Building on the characteristic of profile-based methods, Phyling incorporates a checkpoint mechanism in its align module. This feature allows users to adjust the sample set for phylogenetic analysis while reusing previous search results. When a search is completed, the results are saved to a checkpoint file, which can be loaded during a rerun. This ensures that only newly added or modified samples are searched, greatly reducing computation time for large datasets. It’s important to note that the checkpoint functionality is limited to the search step, as the outputs it generates differ fundamentally and cannot be reused in subsequent steps.

Beyond reruns, the checkpoint system in Phyling also serves as a powerful feature for extending analyses. Hits are associated with samples using a hash of their file content, rather than relying on file names or paths. This content-based identification makes checkpoint files portable and shareable. Along with the original sequencing files, an existing checkpoint can serve as a prebuilt backbone to match new samples. This approach enables efficient integration of additional samples into established phylogenies, making it especially useful for identifying and placing newly sequenced or uncharacterized species within a broader evolutionary framework.

In summary, Phyling offers a robust, efficient, and scalable solution for phylogenetic inference in comparative and population genomics. Its ability to integrate well-established marker sets, handle large and evolving datasets, and provide meaningful support metrics makes it a versatile tool for phylogenomic studies. Additionally, the simplified interface lowers the barrier for less experienced users to perform phylogenetic analyses, while advanced users retain the flexibility to customize their workflows using intermediate outputs.

## Availability and implementation

The Phyling package is available at https://github.com/stajichlab/PHYling and Zenodo doi: 10.5281/zenodo.1257001. The analyses described in this study were performed using the release version 2.3.0 archived as doi: 10.5281/zenodo.16415735.

## Acknowledgements

JES was supported by grants from the National Science Foundation (NSF) DEB-1441715, EF-2125066 and subawards of National Institutes of Health (NIH) awards R01AI130128 and R01AI127548. C-H.T and JES were also supported by the U.S. Department of Agriculture, National Institute of Food and Agriculture grant 2020-70029-33202 and Hatch project CA-R-PPA-211-5062-H. JES is a CIFAR Fellow in the program Fungal Kingdom: Threats and Opportunities. Computations were performed using the computer clusters and data storage resources of the HPCC, which were funded by grants from NSF (MRI-2215705, MRI-1429826) and NIH (1S10OD016290-01A1).

