## Supplementary tables and figures for "Phyling: phylogenetic inference from annotated genomes"

| Tool | Module | Dataset | Mean | Standard deviation | Standard error of mean |
|---|---|---|---|---|---|
| Phyling | align | bacterial_distant | 1.95 | 0.01 | 0.00 |
| Phyling | filter | bacterial_distant | 0.32 | 0.00 | 0.00 |
| Phyling | consensus | bacterial_distant | 2.24 | 0.01 | 0.00 |
| Phyling | concat | bacterial_distant | 473.44 | 0.29 | 0.17 |
| Phyling | FT only | bacterial_distant | 8.08 | 0.02 | 0.01 |
| OrthoFinder | search | bacterial_distant | 644.93 | 73.81 | 42.61 |
| OrthoFinder | tree | bacterial_distant | 8.52 | 1.04 | 0.60 |
| GToTree | GToTree | bacterial_distant | 64.77 | 5.08 | 2.93 |
| Phyling | align | fungal_distant | 52.36 | 0.14 | 0.08 |
| Phyling | filter | fungal_distant | 3.46 | 1.11 | 0.64 |
| Phyling | consensus | fungal_distant | 2.91 | 0.05 | 0.03 |
| Phyling | concat | fungal_distant | 586.42 | 13.38 | 7.73 |
| OrthoFinder | search | fungal_distant | 2,684.34 | 1.07 | 0.62 |
| OrthoFinder | tree | fungal_distant | 26.19 | 0.82 | 0.47 |

**Table S1.** Runtime of different tools and modules. The “Dataset” column indicates the dataset used for benchmarking. “Mean”, “Standard deviation”, and “Standard error of mean” are represented in minutes, based on three replicates.

| Tool | Module | Dataset | Mean | Standard deviation | Standard error of mean |
|---|---|---|---|---|---|
| Phyling | align | bacterial_distant | 3.58 | 0.08 | 0.05 |
| Phyling | filter | bacterial_distant | 0.44 | 0.01 | 0.00 |
| Phyling | consensus | bacterial_distant | 0.64 | 0.00 | 0.00 |
| Phyling | concat | bacterial_distant | 5.35 | 0.08 | 0.04 |
| Phyling | FT only | bacterial_distant | 1.04 | 0.00 | 0.00 |
| OrthoFinder | search | bacterial_distant | 36.39 | 0.06 | 0.04 |
| OrthoFinder | tree | bacterial_distant | 15.92 | 4.98 | 2.87 |
| GToTree | GToTree | bacterial_distant | 4.06 | 0.05 | 0.03 |
| Phyling | align | fungal_distant | 12.51 | 0.84 | 0.49 |
| Phyling | filter | fungal_distant | 0.84 | 0.11 | 0.06 |
| Phyling | consensus | fungal_distant | 1.00 | 0.02 | 0.01 |
| Phyling | concat | fungal_distant | 9.04 | 0.02 | 0.01 |
| OrthoFinder | search | fungal_distant | 110.08 | 1.49 | 0.86 |
| OrthoFinder | tree | fungal_distant | 60.22 | 0.01 | 0.01 |

**Table S2.** Peak memory usage of different tools and modules. The “dataset” column indicates the dataset used for benchmarking. “Mean”, “Standard deviation”, and “Standard error of mean” are represented in GB, based on three replicates.

| | Phyling | | GToTree | | OrthoFinder | |
|---|---|---|---|---|---|---|
| | #Taxa | #Tips | #Taxa | #Tips | #Taxa | #Tips |
| Total | 50 | 251 | 50 | 251 | 50 | 251 |
| Monophyletic | 18 | 171 | 20 | 176 | 18 | 144 |
| Non-Monophyletic | 7 | 54 | 5 | 49 | 7 | 81 |
| Monotypic | 25 | 25 | 25 | 25 | 25 | 25 |
| Intruder | 3 | 4 | 4 | 5 | 17 | 32 |
| Outlier | 4 | 4 | 2 | 2 | 3 | 4 |

**Table S3.** Monophyly test summary of phylogenetic inference made by each tool using the bacterial distant dataset. The “#Taxa” column indicates the number of phylum-level taxa assigned to each category, and the “#Tips” column represents the total number of tips falling into each category. Row “Monotypic” refers to taxa represented by only a single tip; “Intruder” refers to clades that appear within another taxon’s clades; “Outliers” indicates single-tip intruders which are too far away from the rest of their taxon.

### Supplementary figures

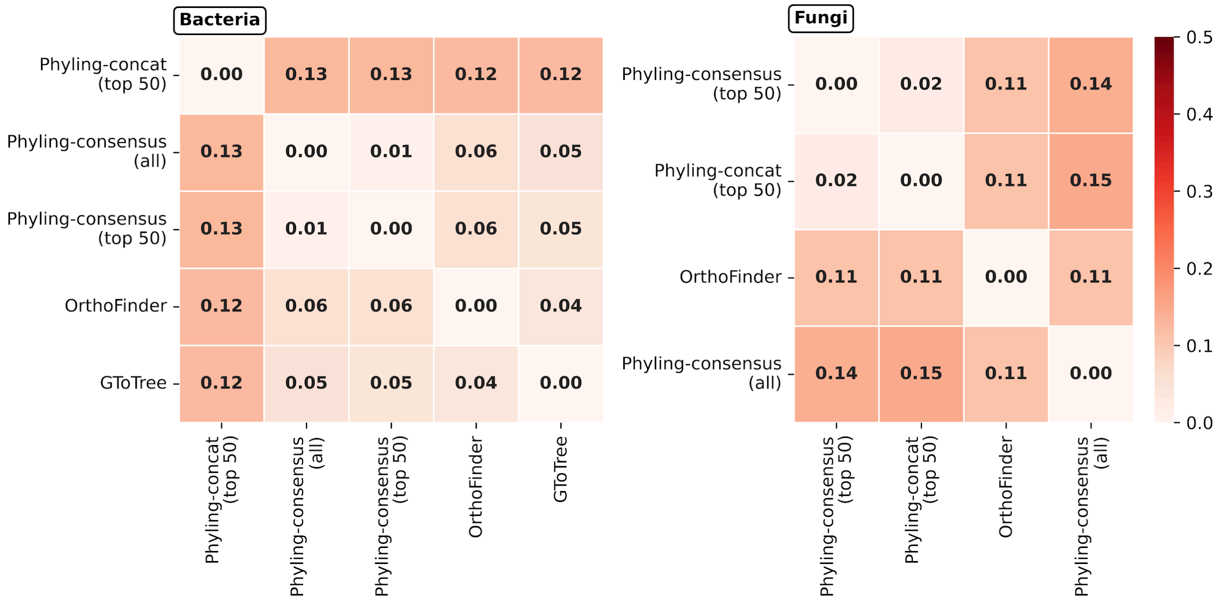


**Figure S1.** Generalized RF distance matrices among trees inferred from general datasets.

The left and right matrices represent results from the bacterial and fungal general datasets, respectively. Matrices are hierarchically clustered. While RF distances range from 0 to 1, the color scale is limited to 0 to 0.5 range to highlight differences.

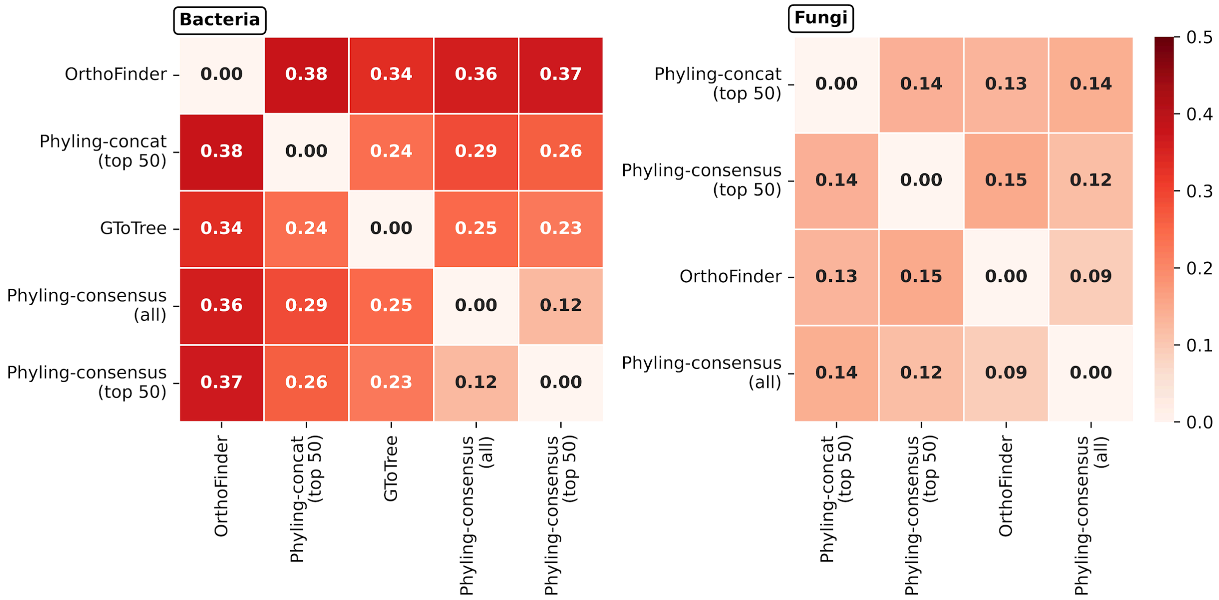


**Figure S2.** Generalized RF distance matrices among trees inferred from distant datasets. The left and right matrices represent results from the bacterial and fungal distant datasets, respectively. Matrices are hierarchically clustered. While RF distances range from 0 to 1, the color scale is limited to 0 to 0.5 range to highlight differences.

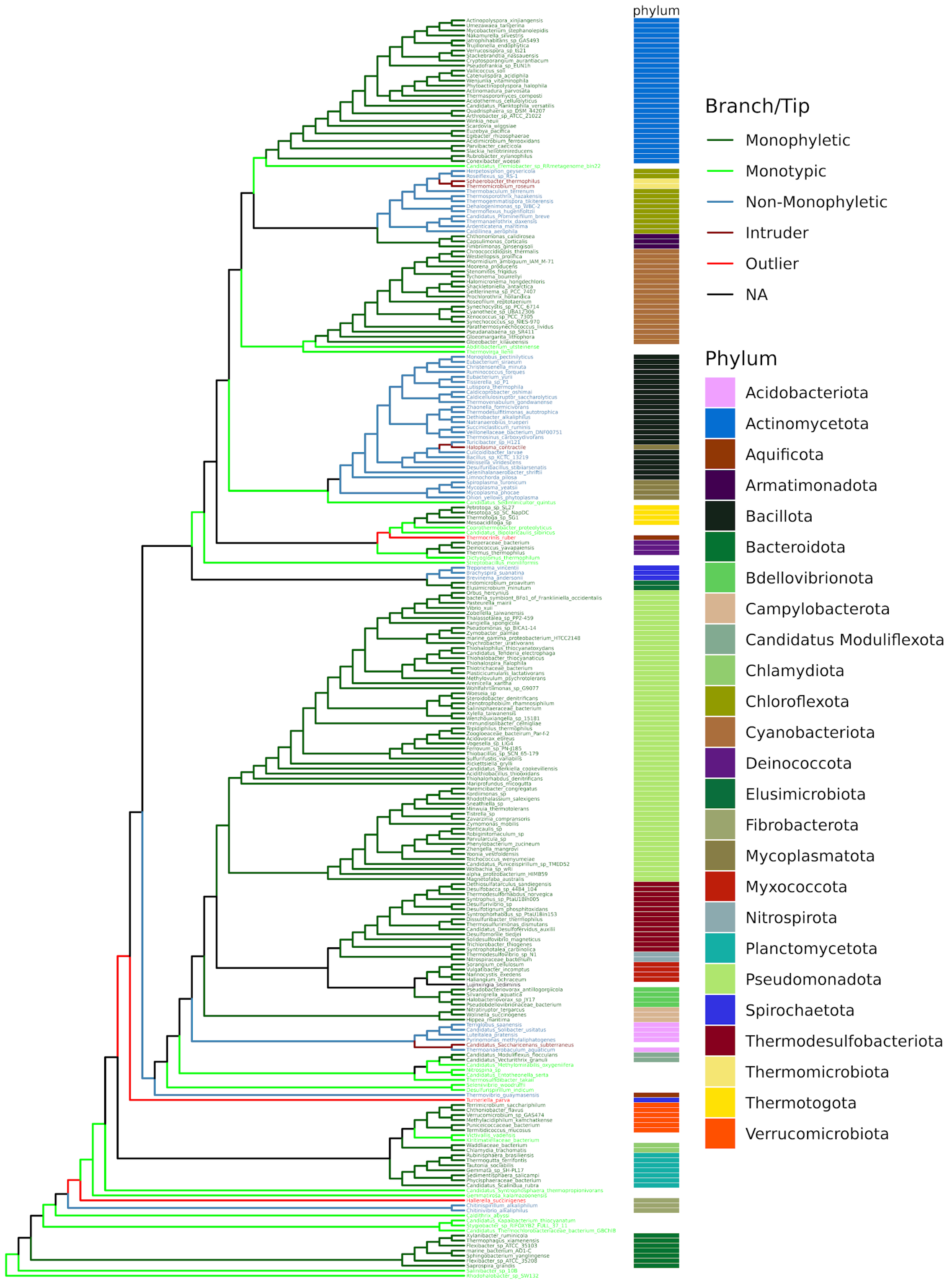


**Figure S3.** Cladogram of the phylogenetic inference made by Phyling consensus mode using the bacterial distant dataset. The branch and tip colors indicate monophyly status, where the blue branches, dark red and red highlight non-monophyletic clades and their corresponding intruding tips, indicate potential misclassifications. Color labels on the right represent the phylum-level taxonomic assignments for each tip, excluding monotypic tips.

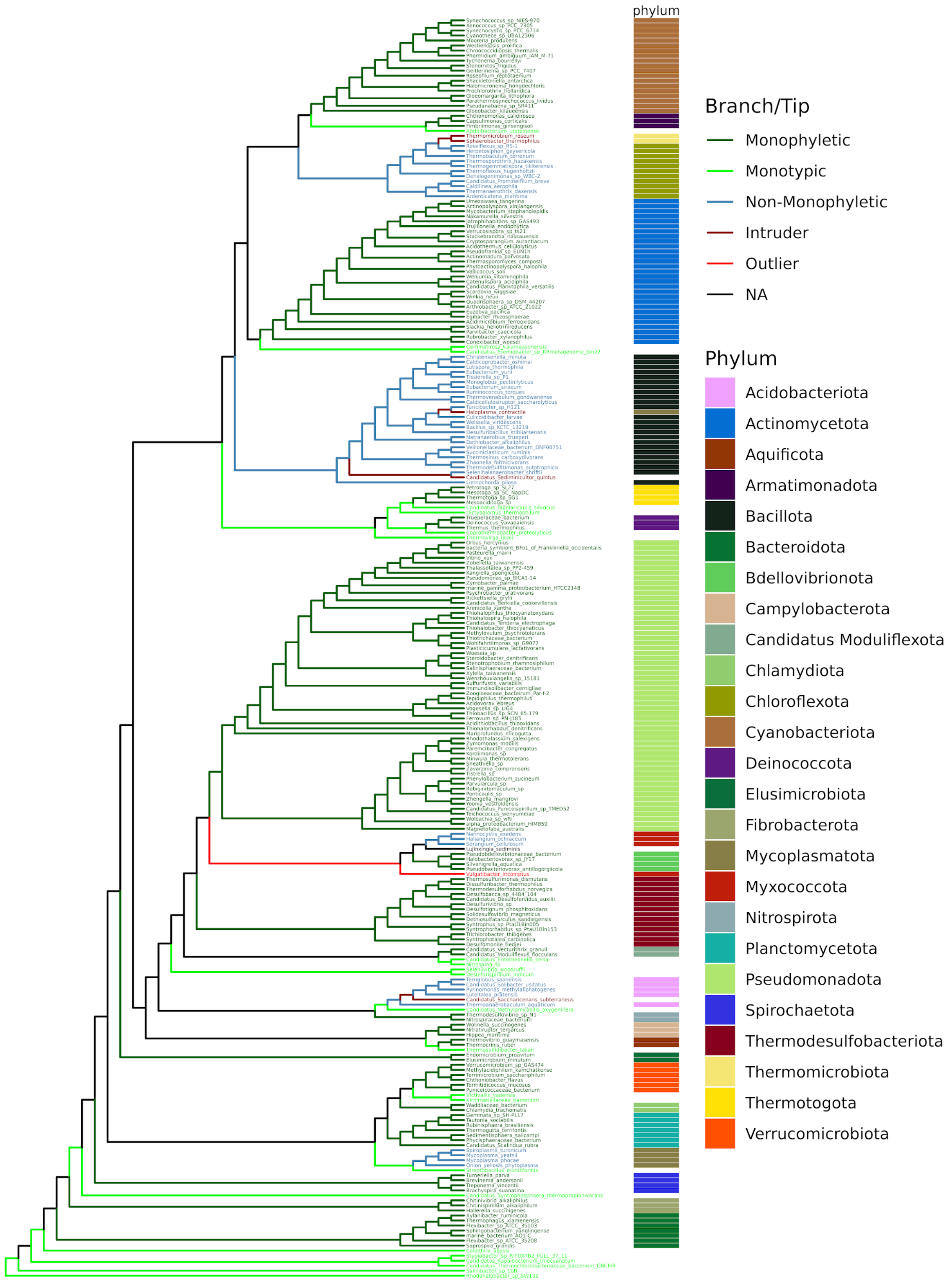


**Figure S4.** Cladogram of the phylogenetic inference made by GToTree using the bacterial distant dataset. Phylum-level taxonomic assignments follow the same color code as Supplementary Fig. 3.

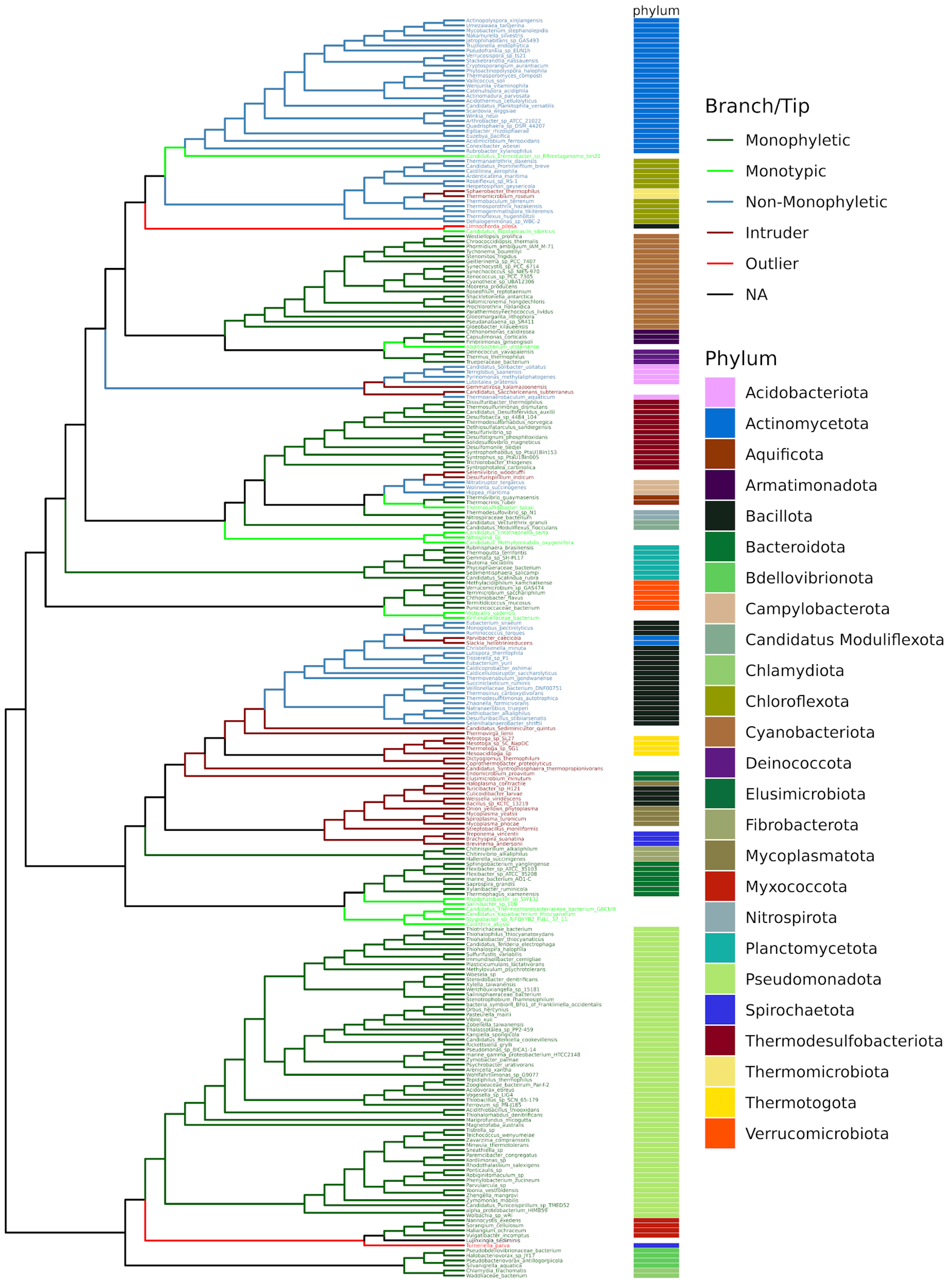


**Figure S5.** Cladogram of the phylogenetic inference made by OrthoFinder using the bacterial distant dataset. Phylum-level taxonomic assignments follow the same color code as Supplementary Fig. 3.

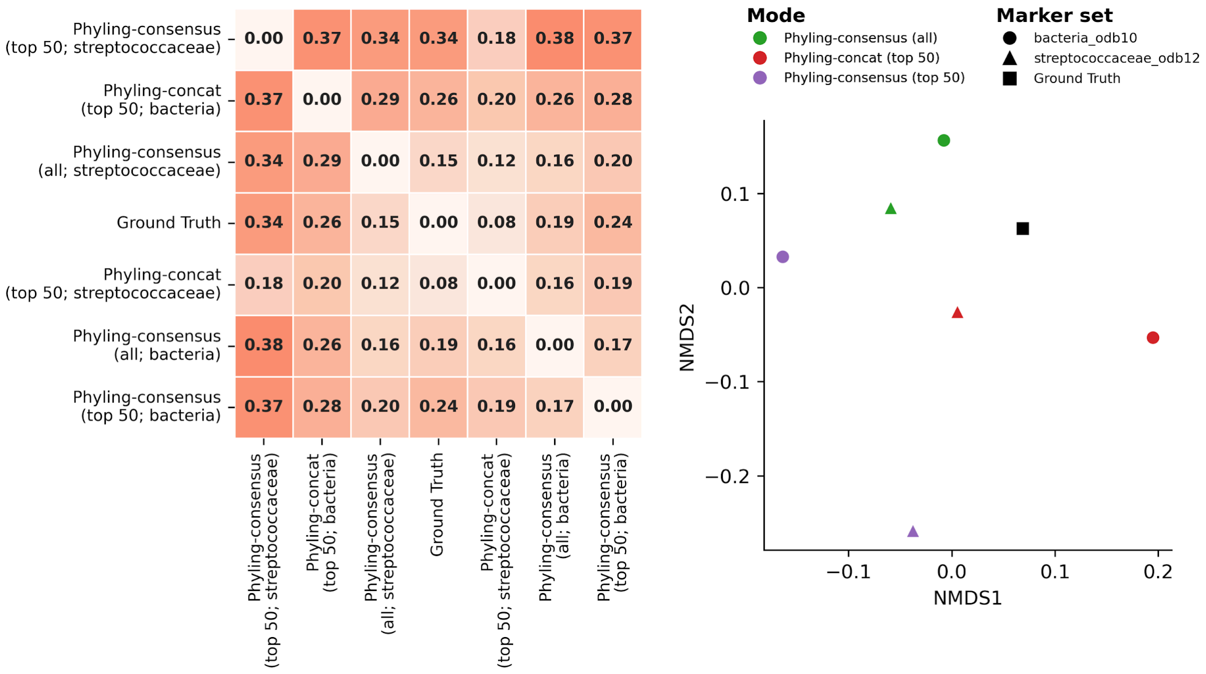


**Figure S6.** Generalized RF distance matrix and its corresponding NMDS plot among trees inferred using orthologs identified by different marker sets. The distance matrix on the left is hierarchical clustered, with values ranging from 0 to 1. In the NMDS plot, colors represent different modes, shapes represent marker sets, and the black square indicates the ground truth. All inferences are made using protein-coding nucleotide sequences.
